# *Chlamydia muridarum* Infection Impacts Murine Models of Intestinal Inflammation and Cancer

**DOI:** 10.1101/2025.05.14.654029

**Authors:** Glory Leung, Sebastian E. Carrasco, Anthony J Mourino, Mert Aydin, Renata Mammone, Gregory F. Sonnenberg, Gretchen Diehl, David Artis, Hiroshi Yano, Rodolfo J Ricart Arbona, Neil S Lipman

**Author notes:** Authors contributed equally (Rodolfo J Ricart Arbona, Neil S Lipman). Animal Care and Veterinary Service, University of Ottawa, Ottawa, ON.

## Abstract

*Chlamydia muridarum* (Cm) has reemerged as a prevalent infectious agent in research mouse colonies. Despite its prevalence and ability to persistently colonize the murine gastrointestinal tract, few studies have evaluated the potential impact of Cm on experimental models of gastrointestinal disease. Studies were conducted to evaluate the impact of Cm on the *Citrobacter rodentium* (Cr), *Trichuris muris* (Tm) and *Il*10^-/-^ mouse models of intestinal inflammation, as well as on tumorigenesis in the Apc^Min/+^ mouse following administration of DSS. Naïve C57BL/6J (B6), B6.129P2-*Il*10*^tm1Cgn^*/J (*Il*10^-/-^) and C57BL/6J-Apc^Min^/J (Apc^Min/+^) mice were infected with Cm by cohousing with chronic Cm-shedding BALB/cJ mice for 2 weeks; controls were cohoused with Cm-free mice. After cohousing, B6 mice (n=8 Cm-infected and free) were infected with *Citrobacter rodentium* (Cr; 10^9^ CFU orally) or *Trichuris muris* (Tm; 200 ova orally). *Il1*0^-/-^ mice (n=8/group with and without *Helicobacter hepaticus* [10^8^ CFU/mouse] and with and without Cm) and Apc^Min/+^ mice (n=8/group) received 2% DSS for 7 days in drinking water after cohousing. Mice were sacrificed 14-days post-Cr infection, 18-days post-Tm infection, 70-days after cessation of cohousing with *Il*10^-/-^ mice, and 28-days post-DSS administration to Apc^Min/+^ mice. The severity of the cecal and colonic lesions were evaluated and graded using a tiered, semi-quantitative scoring system. Cm infection attenuated colitis associated with Cr (p=0.03), had no effect on Tm associated pathology (p=0.22), worsened colitis in *Il*10^-/-^ mice in the absence of *H. hepaticus* (p=0.007), and reduced chemically induced colonic tumorigenesis in Apc^Min/+^ mice (p=0.004). Thus, Cm colonization differentially impacts several models of intestinal inflammation and tumorigenesis, and the presence of this bacterium in mouse colonies should be considered as a variable in these experimental readouts.

## Introduction

*Chlamydia muridarum* (Cm), a Gram negative obligate intracellular bacterium, was initially described in the 1940’s as mouse pneumonitis virus in mice used to investigate the causes of human respiratory infections.^1^ Approximately a decade later the organism was identified as *Chlamydia trachomatis* and in the 1990s was reclassified as Cm, the only *Chlamydia* species known to naturally infect the family Muridae. Cm had not been detected in research mice since these original descriptions, although it has been used extensively to model genital tract infections in humans caused by *C. trachomatis*.^2^

We recently reported on the unexpected reemergence of Cm in research mouse colonies. About a third of the mice imported to our institution, principally from other U.S. and European academic institutions, were infected with Cm.^1^ Surveys of approximately 900 and 11,000 murine diagnostic and environmental samples submitted to research animal diagnostic laboratories demonstrated that 16% and 14%, respectively, were qPCR-positive for Cm.^1^ This prevalence is markedly higher than most excluded murine adventitious agents including MHV and MPV which were recently reported to have less than a 0.3% prevalence.^3^

Cm has a two-stage life cycle consisting of the infectious, nonreplicating elementary body (EB) and a replicating, non-infectious reticulate body (RB).^2^ Natural transmission is primarily fecal-oral.^4^ Natural infection of immunocompetent mouse strains is not associated with clinical signs, while immunocompromised strains develop respiratory signs, weight loss and pulmonary pathology following natural infection.^4–6^ In immunocompetent mice, Cm elicits IgG secretion and transient lymphocyte proliferation in draining mesenteric lymph nodes as well as induction of monocytes and activation of T-cell subsets resulting in a significant reduction of Cm within the gut, but the bacterium persists in the cecum and large intestine while being shed chronically in the feces.^4,6^

Despite the high prevalence, little is known about the potential confounding effects that Cm infection may have on research models. Since most mouse strains are persistently infected with Cm, we speculated that Cm may impact murine research models used to study intestinal inflammation and neoplasia. Wang *et al*. recently showed that oral inoculation of Cm protects against DSS-induced colitis, likely by promoting IL-22 secretion, a cytokine with strong anti-microbial properties.^7^ The DSS-induced model of colitis is only one of many commonly utilized murine models of intestinal inflammation that could potentially be impacted by Cm. Based on the pathways involved in the pathophysiology of the associated disease, we investigated the impact of Cm on the *Citrobacter rodentium* (Cm)*, Trichuris muris* (Tm) and IL-10 knock out models of intestinal inflammation as well as in the Apc^Min/+^ mouse model of tumorigenesis.

## Materials and Methods

### Experimental Models

To investigate the impact of Cm infection on various murine gastrointestinal models, we first infected C57BL/6J, B6.129P2-Il10*^tm1Cgn^*/J (*Il*10^-/-^) and C57BL/6J-Apc^Min^/J (Apc^Min/+^) mice with Cm by cohousing with chronically shedding Cm-infected BALB/cJ (C) mice for 2-weeks (1 infected C mouse/1-4 naïve experimental mice). Naïve Cm-free C mice were cohoused with the same strains as controls. The experimental design and timeline are depicted graphically in Figure 1.

**Figure 1:**
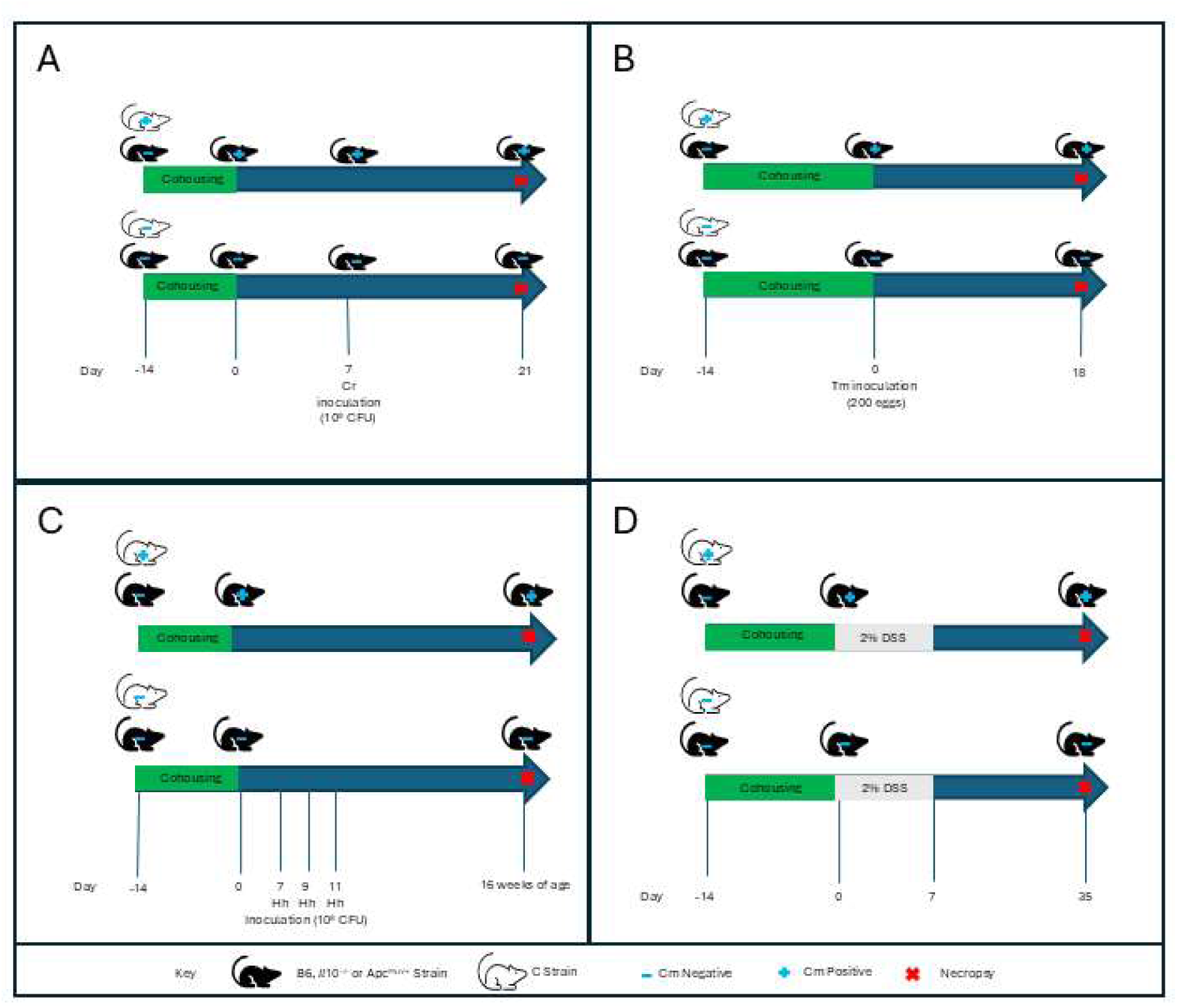
Graphical representation of the study design and timeline. Experimental design and timeline for (A) *Citrobacter rodentium,* (B) *Trichuris muris*, (C) *Il*10^-/-^, and (D) Apc^Min/+^ experiments. *Citrobacter rodentium* (*Cr*), *Trichuris muris* (Tm) and *Helicobacter hepaticus* (Hh) were administered via gavage. DSS was provided in the drinking water.

Cm-positive and -negative B6 mice, confirmed by qPCR, were inoculated with either *Citrobacter rodentium* (Cr; n=8/Cm+ or Cm-; 10^9^/CFUs) or *Trichuris muris* (Tm; n=8/Cm+ or Cm-; 200 ova) via orogastric gavage 14-days after cohousing with Cm-shedding or Cm-free C mice. Fourteen or 18-days post-Cr or -Tm infection, respectively, mice were euthanized, feces collected for Cm and Cr qPCR (Cr-infected mice only), a complete necropsy was performed and the intestinal lesions were scored using a modified published system.^8,9^

*Il*10^-/-^ mice (n=32), 16 of which had been cohoused with Cm-shedding and the remaining with Cm-free C mice, were inoculated with *H. hepaticus* (Hh; 10^8^ CFUs; n=16) or saline (n=16) via orogastric gavage every 2 days for a total of 3 doses at 7 weeks of age. At 16 weeks of age, mice were euthanized, feces collected for Cm and Hh qPCR, a complete necropsy was performed, and the intestinal lesions were scored using a modified scoring system.^10^

Apc^Min/+^ mice (n=8/8; Cm-infected/Cm-free) were provided 2% dextran sodium sulphate (DSS; Sigma-Aldrich, St. Louis, MO) in drinking water for 7 days immediately after cohousing with Cm-infected or Cm-free C mice for 2 weeks. Four weeks post-DSS treatment, mice were euthanized and the number and location of gross tumors in the gastrointestinal tract (GIT) were counted under a dissecting microscope. Select tumors from each animal were analyzed histologically to confirm malignancy.

### Animals

Female C57BL/6J (B6; n=32; 6-week-old; The Jackson Laboratory, Bar Harbor, ME) mice were used for *Citrobacter rodentium* and *Trichuris muris* experiments (n=16/experiment); female B6.129P2-Il10*^tm1Cgn^*/J (*Il*10^-/-^; n=32; 4-week-old; The Jackson Laboratory) were used to assess inflammatory bowel disease; and, female C57BL/6J-ApcMin/J (Apc^Min/+^; n=16; 4-week-old; The Jackson Laboratory) were used to assess tumorigenesis. Female Cm-infected and Cm-free BALB/cJ (C; n=4 and n=3, respectively; >16 weeks old; The Jackson Laboratory, Bar Harbor, ME) mice were used for cohousing.

On study initiation all mice were free of ectromelia virus, Theiler meningoencephalitis virus (TMEV), Hantaan virus, K virus, LDH elevating virus (LDEV), lymphocytic choriomeningitis virus (LCMV), mouse adenovirus (MAV), murine cytomegalovirus (MCMV), murine chapparvovirus (MuCPV), mouse hepatitis virus (MHV), minute virus of mice (MVM), murine norovirus (MNV), mouse parvovirus (MPV), mouse thymic virus (MTV), pneumonia virus of mice (PVM), polyoma virus, reovirus type 3, epizootic diarrhea of infant mice (mouse rotavirus, EDIM), Sendai virus, murine astrovirus-2 (MuAstV-2), Bordetella spp., *Citrobacter rodentium, Chlamydia muridarum, Clostridium piliforme, Corynebacterium bovis, Corynebacterium kutscheri, Filobacterium rodentium* (CAR bacillus), *Mycoplasma pulmonis*and other *Mycoplasma* spp., *Salmonella* spp., *Streptobacillus moniliformis, Helicobacter* spp., *Klebsiella pneumoniae* and *K. oxytoca, Pasteurella multocida, Rodentibacter pneumotropicus/heylii, Pseudomonas aeruginosa, Staphylococcus aureus, Streptococcus pneumoniae* and Beta-hemolytic *Streptococcus* spp., *Yersinia enterocolitica* and *Y. pseudotuberculosis*, *Proteus mirabilis, Pneumocystis murina*, *Encephalitozoon cuniculi*, ectoparasites (fleas, lice, and mites), endoparasites (tapeworms, pinworms, and other helminths), protozoa (including *Giardia* spp. and *Spironucleus* spp.), *Toxoplasma gondii,* trichomonads, and dematophytes.

### Husbandry and Housing

Mice were maintained in autoclaved, individually ventilated, polysulfone shoebox cages with stainless-steel wire-bar lids and filter tops (IVC; no. 19, Thoren Caging Systems, Inc., Hazelton, PA) on autoclaved aspen chip bedding (PWI Industries, Quebec, Canada) at a density of no greater than 5 mice per cage. Each cage was provided with a Glatfelter paper bag containing 6 g of crinkled paper strips (EnviroPak, WF Fisher and Son, Branchburg, NJ) and a 2-inch square of pulped virgin cotton fiber (Nestlet, Ancare, Bellmore, NY) for enrichment. Mice were fed a natural ingredient, closed source, γ-irradiated, autoclaved diet (LabDiet 5KA1, PMI, St. Louis, MO) ad libitum.^11^ All animals were provided autoclaved reverse osmosis acidified (pH 2.5 to 2.8 with hydrochloric acid) water in polyphenylsulfone bottles with stainless-steel caps and sipper tubes (Techniplast, West Chester, PA). Cages were changed every 7 days within a class II, type A2 biological safety cabinet (LabGard S602-500, Nuaire, Plymouth, MN). All cages were housed within a dedicated, restricted-access cubicle, which was maintained on a 12:12-h light:dark cycle (on 6 AM; off 6 PM), relative humidity of 30 to 70%, and room temperature of 72 ± 2°F (22.2 ± 1.1°C). All cage changing and animal handling was performed by the author (GL). The animal care and use program at Memorial Sloan Kettering Cancer Center (MSK) is accredited by AAALAC International and all animals are maintained in accordance with the recommendations provided in the Guide.^12^ All animal use described in this investigation was approved by MSK’s IACUC in agreement with AALAS’ position statements on the Humane Care and Use of Laboratory Animals and Alleviating Pain and Distress in Laboratory Animals.^12^

### Infection of Experimental Mice with Cm via Cohousing

The generation of chronically Cm-infected C mice was previously described.^6, 1^ Briefly, Cm qPCR negative C mice were inoculated with 2.72×10^3^ IFU of a previously isolated Cm field strain in 100μL sucrose-phosphate-glutamic acid buffer (SPG, pH 7.2) via orogastric gavage. A new, sterilized gavage needle (22g × 38.1mm, Cadence Science, Cranston, RI) was used for each cage.

### Citrobacter rodentium (Cr), Trichuris muris (Tm) and Helicobacter hepaticus (Hh) Culture and Inoculation

Cr (Cat#51549, ATCC, Manassus, VA) was cultured in Luria-Bertani broth overnight (Cat# 3002121, MP Biomedicals, Santa Ana, CA) at 37°C with constant shaking. The administered inoculum was confirmed by plating bacteria on MacConkey agar (Cat# 212123, BD Diagnostics, Franklin Lakes, NJ) and counting the colonies after 14-18 hours of incubation. B6 mice were inoculated with 1×10^9^ CFUs in 200 μL of PBS via orogastric gavage. A new, sterilized gavage needle (24g x 38.1mm, Cadence Science) was used for each cage.

Embryonated Tm eggs (Artis Laboratory, Weill Cornell Medicine, NY, NY) were cultivated and maintained in the dark at 4°C, and the inoculum prepared as previously described.^13^ Briefly, Tm was maintained in Rag1^-/-^ animals. Following sacrifice, adult nematodes were isolated from the cecum and proximal colon and cultured at 37°C in serum-free Roswell Park Memorial Institute Medium 1640 (Corning Inc., Corning, NY) containing 500 U/ml penicillin and 500 μg/ml streptomycin. Eggs were collected from the culture by filtering through a 70 µm or 100 µm cell strainer, washed thrice in sterile water, and incubated at room temperature in the dark in a ventilated T75 cell culture flask (Corning Inc.) for 6-8 weeks. Embryonation was confirmed under a dissection microscope, and the eggs were stored at 4°C in the dark after adjusting the concentration to 200 embryonated eggs per 200 µL in water. B6 mice were inoculated with 200 embryonated Tm eggs in 200 µL water via oral gavage using a new, sterilized gavage needle (24g x 38.1mm, Cadence Science).

Hh (51449™, ATCC) was cultured on blood agar plates (TSA with 5% sheep blood, Thermo Fisher Inc., Waltham, MA) as previously described.^14^ Inoculated plates were placed into a Billups-Rothenberg hypoxia chamber (Embrient Inc., San Diego CA) and an anaerobic gas mixture consisting of 80% nitrogen, 10% hydrogen, and 10% carbon dioxide (Airgas, New York, NY) was added to create a micro-aerophilic atmosphere in which the oxygen concentration was 3 - 5%. The concentration of bacterial inoculation dose was determined using a spectrophotometric optical density (OD) analysis at 600 nm and adjusted to OD600 readings between 1 and 1.5. Aliquots were then frozen at -80°C in Brucella broth with 20% glycerol. Frozen aliquots were thawed on ice prior to oral inoculation. *Il*10^-/-^ mice were inoculated with 10^8^ CFUs of Hh in 0.2ml of culture broth via orogastric gavage every 2 days for a total of 3 doses. A new, sterilized gavage needle (24g x 38.1mm, Cadence Science) was used for each cage.

### Clinical Monitoring

All mice were monitored daily to assess activity, respiratory rate and effort, coat condition and posture, signs of rectal prolapse or diarrhea, and body condition score. Weights were recorded at the start of each experiment and as needed thereafter based on the BCS. Animals displaying weight loss (20% reduction from baseline) and/or reduced body condition (BCS < 2/5), presenting with dyspnea and/or cyanosis, and/or failing to respond to stimulation were euthanized.

### Fecal Collection

Fecal pellets for PCR were collected ante- or post-mortem and pooled by cage for submission. When collected antemortem, the mouse was first lifted by the base of the tail and allowed to grasp onto the wire-bar lid while a 1.5ml microcentrifuge tube was placed underneath the anus for fecal collection directly into the tube. If the animal did not defecate within 30 seconds, a fecal sample from the cage floor was collected. Post-mortem fecal collection was collected directly from the rectum, if available, or the cage floor.

### Chlamydia muridarum (Cm), Citrobacter rodentium (Cr), and Helicobacter spp. qPCR Assays

DNA and RNA were copurified from fecal samples using the Qiagen DNeasy 96 blood and tissue kit (Qiagen, Hilden, Germany). Nucleic acid extraction was performed using the manufacturer’s recommended protocol, “Purification of Total DNA from Animal Tissues”, with the following buffer volume modifications: 300µL of Buffer ATL + Proteinase K, 600µL of Buffer AL + EtOH, and 600µL of lysate were added to individual wells of the extraction plate. Washes were performed with 600µL of Buffers AW1 and AW2. Final elution volume was 150µL of Buffer AE.

A probe-based PCR assay for Cm was designed using IDT’s PrimerQuest Tool (IDT, Coralville, IA) based on the 16s rRNA sequence of *Chlamydia muridarum*, Nigg strain (Accession NR_074982.1, located in the National Center for Biotechnology Information database). Primer and Probe sequences generated from the PrimerQuest Tool were checked for specificity using NCBI’s BLAST (Basic Local Alignment Search Tool). The probe was labeled with FAM and quenched with ZEN and Iowa Black FQ (IDT, Coralville, IA). Primer names, followed by associated sequences were as follows: *C. muridarum*_For (GTGATGAAGGCTCTAGGGTTG); *C. muridarum*_Rev (GAGTTAGCCGGTGCTTCTTTA), *C. muridarum*_Probe (TACCCGTTGGATTTGAGCGTACCA).

A probe-based PCR assay for Cr was designed using IDT’s PrimerQuest Tool (IDT, Coralville, IA) based on Cr’s espB gene sequence of Cr, (Accession AF_177537.1, located in the National Center for Biotechnology Information database). Primer and probe sequences generated from the PrimerQuest Tool were checked for specificity using NCBI’s BLAST (Basic Local Alignment Search Tool). The probe was labeled with FAM and quenched with ZEN and Iowa Black FQ (IDT, Coralville, IA). Primer names, followed by associated sequences are as follows: *C. rodentium*_For (CAGGTATCGCTGATGATGTTACT); *C. rodentium*_Rev (CAGATTTGCCTTCCGTGTTAAAT), *C. rodentium*_Probe (TGCTCAGAAAGCTTCTCAGGTAGCTG).

A proprietary probe-based PCR assay for *Helicobacter* spp. was designed by aligning 16s sequences from the following *Helicobacter* spp.: *H. bilis*, *H. ganmani*, *H. hepaticus*, *H. mastromyrinus*, *H. muridarum*, *H. rappini*, *H. rodentium*, *H. trogontum*, and *H. typhlonius*. Sequences were aligned using Sequencher 5.4.6 software (Gene Codes, Ann Arbor, MI). Primers were designed to conserved regions between all species and checked for specificity using NCBI’s BLAST (Basic Local Alignment Search Tool). The probe was labeled with HEX and quenched with ZEN and Iowa Black FQ (IDT, Coralville, IA).

Real time qPCR assays were carried out using a real-time PCR system (BioRad CFX machine, Bio-Rad, Hercules, CA). Reactions were run using Qiagen’s QuantiNova Probe PCR Kit (Qiagen) using the kit’s recommended concentrations and cycling conditions. Final Concentration: 1x QuantiNova Master mix, 0.4 µM Primers, and 0.2 µM FAM or HEX labeled Probe. Cycling: 95°C 2 min, followed by 40 cycles of 95°C 5 sec, 60°C 30 sec.

All reactions were run in duplicate by loading 5µl of template DNA to 15µl of the reaction mixture. A positive and negative, no-template control were included in each run. The positive control was a purified PCR amplicon diluted to produce consistent values of Ct 28. A 16s universal bacterial PCR assay using primers 27_Forward and 1492_Reverse was run on all samples to check for DNA extraction and inhibitors. Samples were considered positive if both replicates had similar values and produced a Ct value of less than 40. A sample was called negative if no amplification from the qPCR assay was detected and a positive amplification from the 16s assay was detected.

### Anatomic Pathology, Lesion Scoring and Tm Infection Confirmation

Following euthanasia by carbon dioxide overdose, a complete necropsy was performed and gross lesions recorded. For all mice except for the Apc^Min/+^ mice, the GIT from the cecum to rectum was removed and placed in cassettes in a Swiss roll configuration and fixed in 10% neutral buffered formalin for at least 72 hours. Tissues were then processed in ethanol and xylene and embedded in paraffin in a tissue processor (Leica ASP6025, Leica Biosystems, Deer Park, IL). Paraffin blocks were sectioned at 5μm, stained with hematoxylin and eosin (H&E), and evaluated blindly by a board-certified veterinary pathologist (SC). The cecum and colon of Cr- and Tm-inoculated mice, and *Il*10^-/-^ mice were graded as described in tables 1 - 2 using a modified scoring system for each model.^8,9,10^ A composite pathology score was calculated for each mouse by combining all subcategory scores. Tm infection was confirmed by detecting nematodes histologically.

**Table 1:** Histopathologic scoring scheme for *Citrobacter rodentium* (Cr) and *Trichuris muris* (Tm) infection in the colon and cecum adapted from Bouadoux *et al*. and Kopper *et al*.^8,9^.

| **Criterion** | **0 (Absence)** | **1 (Minimal)** | **2 (Mild)** | **3 (Moderate)** | **4 (Severe)** |
| --- | --- | --- | --- | --- | --- |
| Hyperplasia | None | 1.5x normal crypt height | 2x normal crypt height | 3x normal crypt height | >4x normal crypt height |
| Goblet cell depletion | None | 5% | 5 - 25% | 25 - 60% | >60% |
| Inflammation | None to multifocal rare individual leukocytes scattered in the lamina propria | Multifocal, small discrete clusters of leukocytes in the lamina propria with or without transepithelial leukocyte infiltration | Multifocal, coalescing lamina propria inflammation with or without early submucosal extension | Multifocal, coalescing lamina propria inflammation with multifocal submucosal extension | Multifocal, diffuse transmural inflammation affecting mucosa, submucosa and deeper layers (muscularis or serosa) |
| Epithelial defects | None | Occasional dilated glands with luminal cell debris (crypt abscesses) | Multifocal crypt abscesses and surface epithelial tattering and/or attenuation (intact) | Multifocal erosions or focal ulceration | Multifocal ulcerations |
| Fibrosis* | None | None or focal, rare in lamina propria | Occasional foci in small segments of the LI, fibroplasia (immature fibroblast proliferation) associated with submucosal inflammation | Multifocal, fibroplasia associated with submucosal inflammation | Multifocal, Mature fibrosis with submucosal inflammation and replacement of normal tissue architecture |

\*Minimal, mild and moderate fibrosis for the Cr model was defined as 1, 1-2 and >2 foci respectively. Minimal, mild and moderate fibrosis for the Tm model was defined as 1-2, 2-5 and >5 foci respectively.

**Table 2:** Histopathologic grading scheme for the colon and cecum from *Il*10^-/-^ mice adapted from Rogers and Houghton^10^.

| **Criterion** | **0 (Absence)** | **1 (Minimal)** | **2 (Mild)** | **3 (Moderate)** | **4 (Severe)** |
| --- | --- | --- | --- | --- | --- |
| Inflammation | None to multifocal rare individual leukocytes scattered in the lamina propria | Multifocal, small discrete clusters of leukocytes in the lamina propria | Multifocal, coalescing lamina propria inflammation with or without early submucosal extension | Multifocal, coalescing lamina propria inflammation with multifocal submucosal extension | Multifocal, diffuse transmural inflammation affecting mucosa, submucosa and deeper layers (muscularis or serosa) |
| Hyperplasia | None | 1.5x normal crypt height | 2x normal crypt height | 3x normal crypt height | >4x normal crypt height |
| Goblet cell depletion | None | 5% | 5 - 25% | 25 - 60% | >60% |
| Epithelial defects | None | Occasional dilated glands with luminal cell debris (crypt abscesses), and surface epithelial tattering and/or attenuation (intact) | Multifocal crypt abscesses, herniation, and surface epithelial tattering and/or attenuation (intact) | Same changes as 2 plus multifocal erosions or focal ulceration | Same as # 3 plus multifocal ulcerations |
| Crypt Atrophy | None | 5-25% | 25-50% | 50-75% | >75% |
| Edema | None | Mild segmental expansion of submucosa | Moderate diffuse submucosal +/- mucosal lamina propria expansion | Severe edema of mucosa and submucosa | Transmural; very severe |
| Dysplasia | None | Mild dysplasia characterized by epithelial cell pleomorphism, plump and attenuated forms, aberrant crypt foci, gland malformation with mild splitting, branching, and infolding, rare cystic dilation | Moderate dysplasia characterized by pleomorphism, early cellular and nuclear atypia, mild piling and infolding, occasional cystic dilation, bulging toward muscularis mucosae and projection into lumen, loss of normal glandular, mucous, or goblet cells, = atypical hyperplasia | Gastrointestinal intraepithelial neoplasia (GIN), severe dysplasia confined to mucosa, features as earlier but greater severity, frequent, and sometimes bizarre mitoses, add 0.5: Intramucosal carcinoma (extension of severely dysplastic regions into muscularis mucosae) | Invasive carcinoma: Submucosal invasion, or any demonstrated invasion into blood or lymphatic vessels, regional nodes, or other metastasis |

Tumorigenesis was assessed in the Apc^Min/+^ mice following euthanasia after which the GIT from the duodenum to the rectum was removed and placed in a Petri dish containing 10% buffered formalin. The entire GIT was then flushed with formalin and opened longitudinally with scissors to expose the lumen for tumor enumeration under a dissecting microscope (Olympus SZX16, Tokyo, JAPAN). Select tumors (n=16) were subsequently submitted for histologic assessment as described above to assess for malignancy.

Immunohistochemistry (IHC) – The entire cecum and colon from each Cm-infected colitis model was processed and stained for Chlamydia MOMP using a technique optimized and validated by MSK’s Laboratory of Comparative Pathology as a secondary confirmatory method.^1^ Briefly, formalin-fixed, paraffin-embedded sections were stained using an automated staining platform (Leica Bond RX, Leica Biosystems, Deer Park, IL). Following deparaffinization and heat-induced epitope retrieval in a citrate buffer at pH 6.0, the primary antibody against Chlamydia MOMP (NB100-65054, Novus Biologicals, Centennial, CO) was applied at a dilution of 1:2000. A rabbit anti-goat secondary antibody (Cat. No. BA-5000, Vector Laboratories, Burlingame, CA) and a polymer detection system (DS9800, Novocastra Bond Polymer Refine Detection, Leica Biosystems) was then applied to the tissues. The 3,3’-diaminobenzidine tetrachloride (DAB) was used as the chromogen, and the sections were counterstained with hematoxylin and examined by light microscopy. Small intestine from an NSG mouse infected with Cm strain Nigg was used as a positive control.^5^ Positive MOMP antigen was identified as chromogenic brown immunolabeling under bright field microscopy.

In situ hybridization (ISH) was conducted on the large intestine of a subset of Cm positive *Il*10^-/-^ animals (n=8) in which the MOMP IHC signal was absent. Briefly, the target probe was designed to detect region 581 to 617 of *Chlamydia muridarum* str. Nigg complete sequence, NCBI Reference Sequence NC_002620.2 (1039538-C1; Advanced Cell Diagnostics, Newark, CA). The target probe was validated on reproductive tracts from mice experimentally inoculated with *Chlamydia muridarum* strain Nigg.^5^ Slides were stained on an automated stainer (Leica Bond RX; Leica Biosystems) with RNAscope 2.5 LS Assay Reagent Kit-Red (322150; Advanced Cell Diagnostics) and Bond Polymer Refine Red Detection (DS9390; Leica Biosystems). Control probes detecting a validated positive housekeeping gene (mouse peptidylprolyl isomerase B, Ppib to confirm adequate RNA preservation and detection; 313918; Advanced Cell Diagnostics) and a negative control, Bacillus subtilis dihydrodipicolinate reductase gene (dapB to confirm absence of nonspecific staining; 312038; Advanced Cell Diagnostics) were used. Positive RNA hybridization was identified as discrete, punctate chromogenic red dots under bright field microscopy.

### Statistical Analysis

Mann-Whitney U (Wilcoxon Rank Sum) tests were used to compare colitis scores between Cm-infected and Cm-free control mice infected with Cr and Tm. The Kruskal-Wallis test for multiple comparisons was used to compare pathology scores between Cm-infected, *Helicobacter hepaticus*-infected, co-infected, and control *Il*10^-/-^ mice. An unpaired t-test was used to compare gross tumor numbers between Cm-infected and Cm-free Apc^Min/+^ mice. For all tests, statistical significance was set as a p ≤ 0.05. Analysis and graphical representations were performed and created in Prism 9 (version 9.3.0, GraphPad Software, Boston, MA) for Windows.

## Results

### Citrobacter rodentium (Cr) colitis model

Mice were confirmed to be infected with Cr only or Cr and Cm as expected at the experimental endpoint via qPCR and IHC. A single Cm- and Cr-infected mouse was found dead 10 days post-inoculation. Histopathology revealed a moderate to marked, multifocal necroulcerative and suppurative colitis. This mouse was excluded from analysis as the lesions may represent an atypical presentation of Cr infection or may have been caused by an unidentified etiology. All remaining animals remained healthy for the duration of the study. The colons of mice infected with both Cr and Cm had statistically significant lower composite and individual scores for inflammation and goblet cell depletion when compared to colons from Cr inoculated, Cm-free mice (Figure 2). Colitis was characterized in the Cm-free mice as diffuse and moderate to marked, while Cm-positive mice had mild to moderate colitis. Both groups had infiltration of the lamina propria by lymphocytes, plasma cells and histiocytes with submucosal extension (Figure 2B & C).

**Figure 2:**
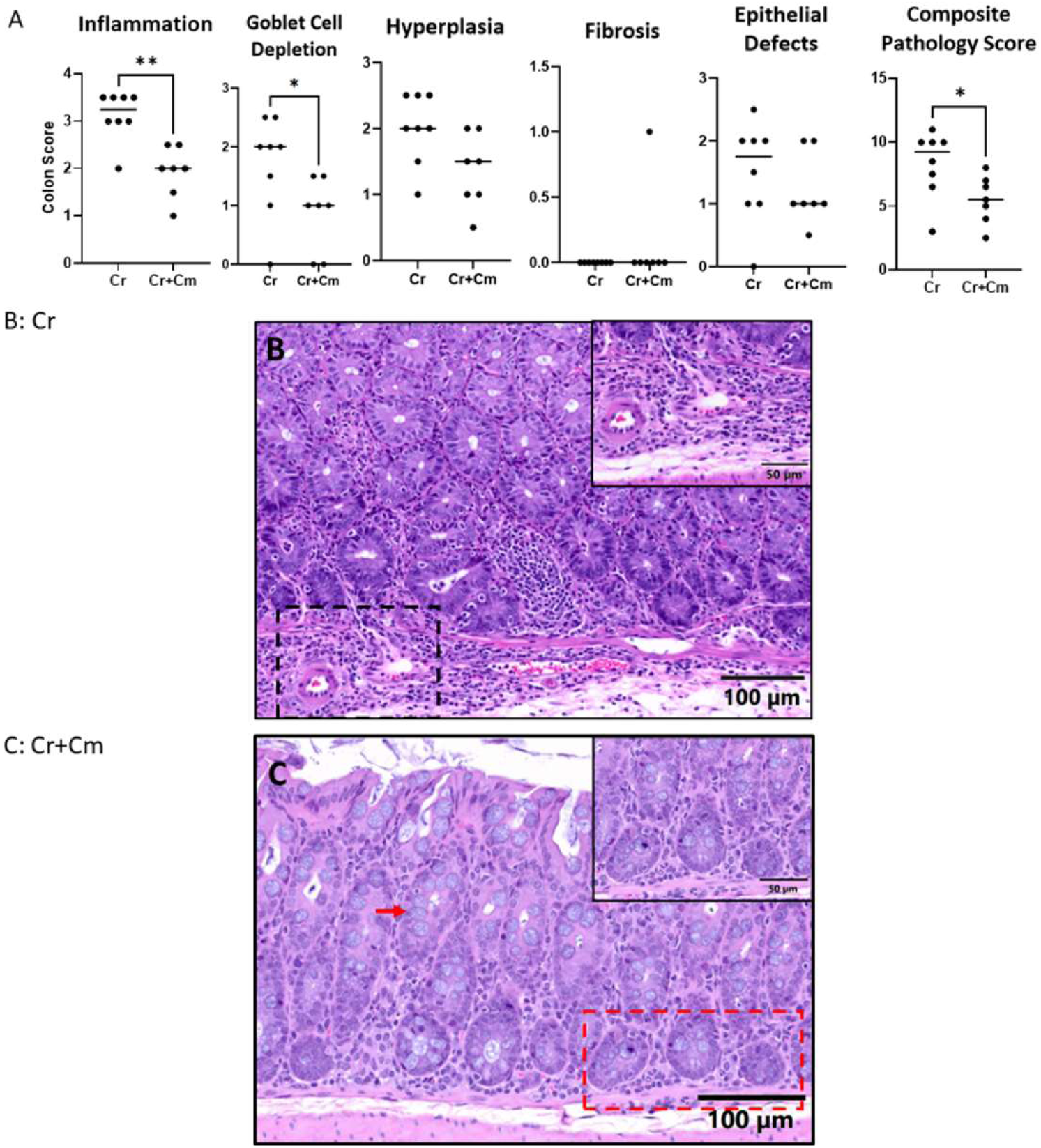
The effects of *Chlamydia muridarum* (Cm) on *Citrobacter rodentium* (Cr)-induced colitis. (A) Individual and composite colitis scores of female C57BL/6J mice infected with only Cr or infected with both Cr and Cm (n=8 mice/group) 14 days post-Cr infection. Composite pathology scores were calculated by combining scores from all subcategories. The crossbar represents the median lesion score for each parameter. (B and C) Representative H&E-stained images of colons from mice infected with only Cr (B) or with both Cr and Cm (C) 14 days post- Cr infection. (B) Moderate to marked mucosal and submucosal inflammation with infiltration of the lamina propria (inset), moderate crypt hyperplasia, and mild goblet cell depletion in affected crypts in the colon of a Cr-infected, Cm-free mice with a composite pathology score of 10 (10x magnification; inset, 40x magnification). (C) Less severe mucosal inflammation (inset), reduced crypt hyperplasia, and less goblet cell depletion (red arrow) in the colon of a Cr-infected, Cm-positive mouse with a composite pathology score of 7 (10x magnification; inset, 40x magnification). *, P≤0.05, **, P≤0.01

### *Trichuris muris* (Tm) colitis model

All mice in each group were confirmed to be infected with only Tm or both Tm and Cm at the end of the experiment by histology and qPCR. No mice developed signs of illness for the duration of the experiment. Mice coinfected with Tm and Cm showed no significant score differences in the cecum and colon for any individual component or the composite score when compared to mice infected only with Tm (Table 3). Colitis in both groups was characterized as diffuse and moderate with infiltration of the lamina propria by lymphocytes, plasma cells, eosinophils and histiocytes with submucosal extension. Tm larvae were readily appreciated in the lumen of all mice examined (Figure 3).

**Figure 3:**
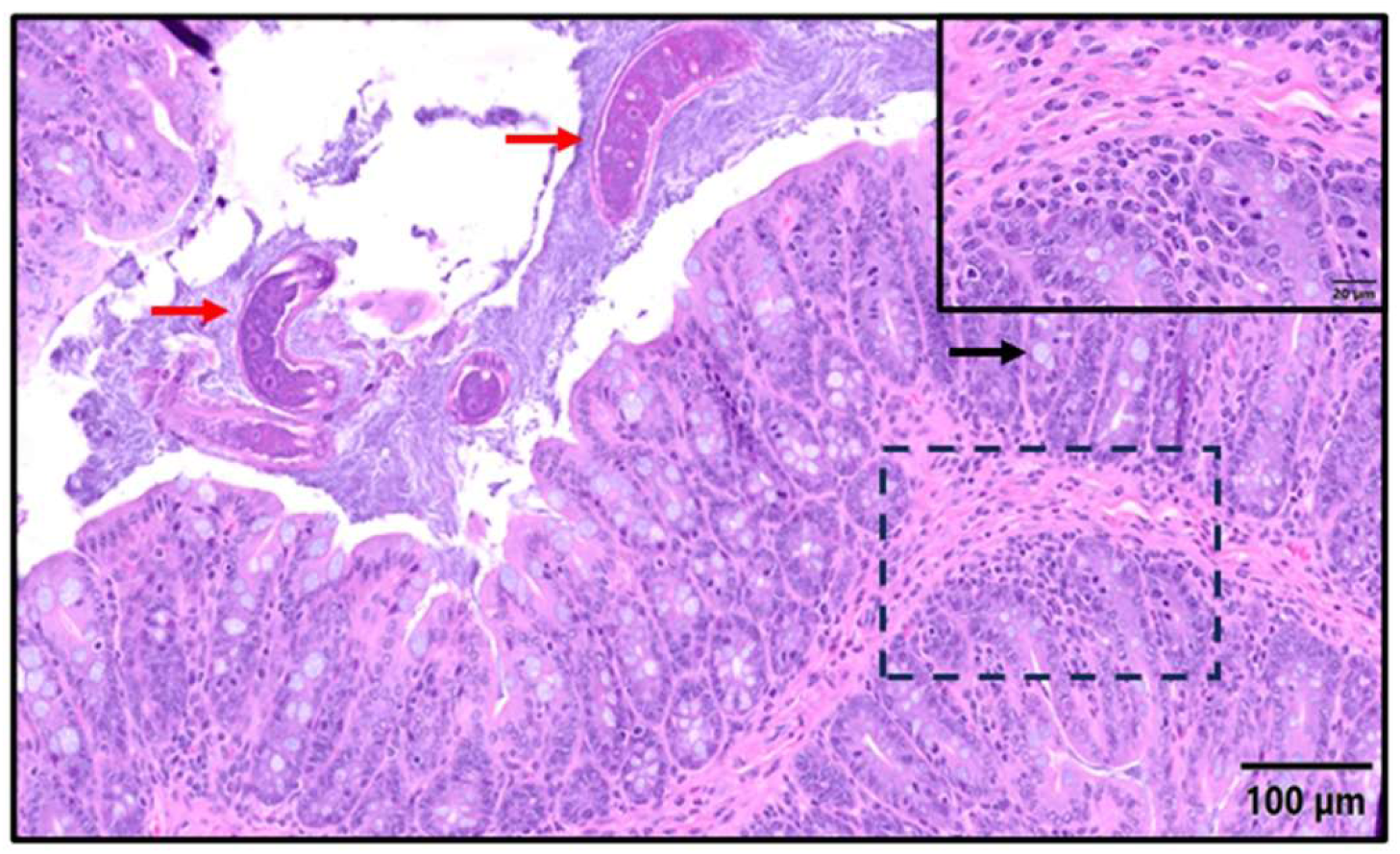
Histopathology of the colon from a *Chlamydia muridarum* (Cm)-free mouse infected with *Trichuris muris* (Tm) 18 days post-Tm infection. Representative H&E-stained image of the colon from a mouse infected with Tm 18 days following cohousing with a Cm-free mouse for 2 weeks (composite pathology score = 7.5). A multifocal infiltrate of lymphocytes, eosinophils and plasma cells is present in the lamina propria with submucosal extension (Inflammation score = 3; inset) in association with mild to moderate hyperplasia and mild goblet cell hypertrophy (goblet cell hypertrophy score = 1; black arrow) in affected crypts and intraluminal Tm larvae (red arrows). (10x magnification; inset, 60x magnification).

**Table 3:** Pathology scores assigned to the cecum and colon of mice infected with *Trichuris muris* (Tm) with and without Chlamydia muridarum *(Cm)*

|  | **Cecum** |  | **Colon** |  |
| --- | --- | --- | --- | --- |
| **Category** | **Tm** | **Cm + Tm** | **Tm** | **Cm + Tm** |
| **Inflammation** | 2.56±0.50 | 2.31±0.46 | 2.25±0.46 | 2.56±0.32 |
| **Epithelial Hyperplasia** | 1.81±0.26 | 1.75±0.46 | 1.5±0.38 | 1.75±0.38 |
| **Goblet Cell Hyperplasia** | 1.25±0.27 | 1.0±0.53 | 1.38±0.35 | 1.19±0.26 |
| **Epithelial Defects** | 1.81±0.46 | 1.5±0.80 | 1.31±0.46 | 1.69±0.37 |
| **Fibrosis** | 0.25±0.46 | 0.44±0.62 | 0.25±0.46 | 0.13±0.35 |
| **Composite Pathology Score** | 7.69±1.56 | 7.0±2.55 | 6.69±1.89 | 7.31±0.88 |

Pathology scores (mean +/- SD) of all criteria of the cecum and colon of female C57BL/6J mice infected with Tm or co-infected with Tm and Cm (n=8 mice/group) 18 days post-Tm infection following cohousing with a Cm-shedding or Cm-free mouse for 2 weeks. None of the score differences in any category nor the composite scores in either the cecum or colon were significantly different.

### Il10^-/-^ colitis model (Helicobacter hepaticus [Hh] infected and free)

Mice were confirmed Cm-free or Cm-infected at euthanasia by qPCR as expected except 2 animals representing a single mouse in the Cm-only group and another in the Cm+Hh coinfected group despite having been qPCR positive after cohousing. Given the initial positive qPCR results after cohousing, and similar intestinal pathology to other mice in the group, these mice were included in analysis. None of the mice developed signs of illness for the duration of the experiment. MOMP staining of the cecum and colon were negative for *Il*10^-/-^ mice cohoused with a Cm-shedding mouse for 2 weeks despite the fact most of these mice were qPCR positive. ISH was also performed on cecum and colon sections to confirm the presence of Cm. Staining was detected in the colon of all Cm-infected animals, except for 1 of the 2 aforementioned qPCR negative mice representing the Cm+Hh coinfected group (Figure 4).

**Figure 4:**
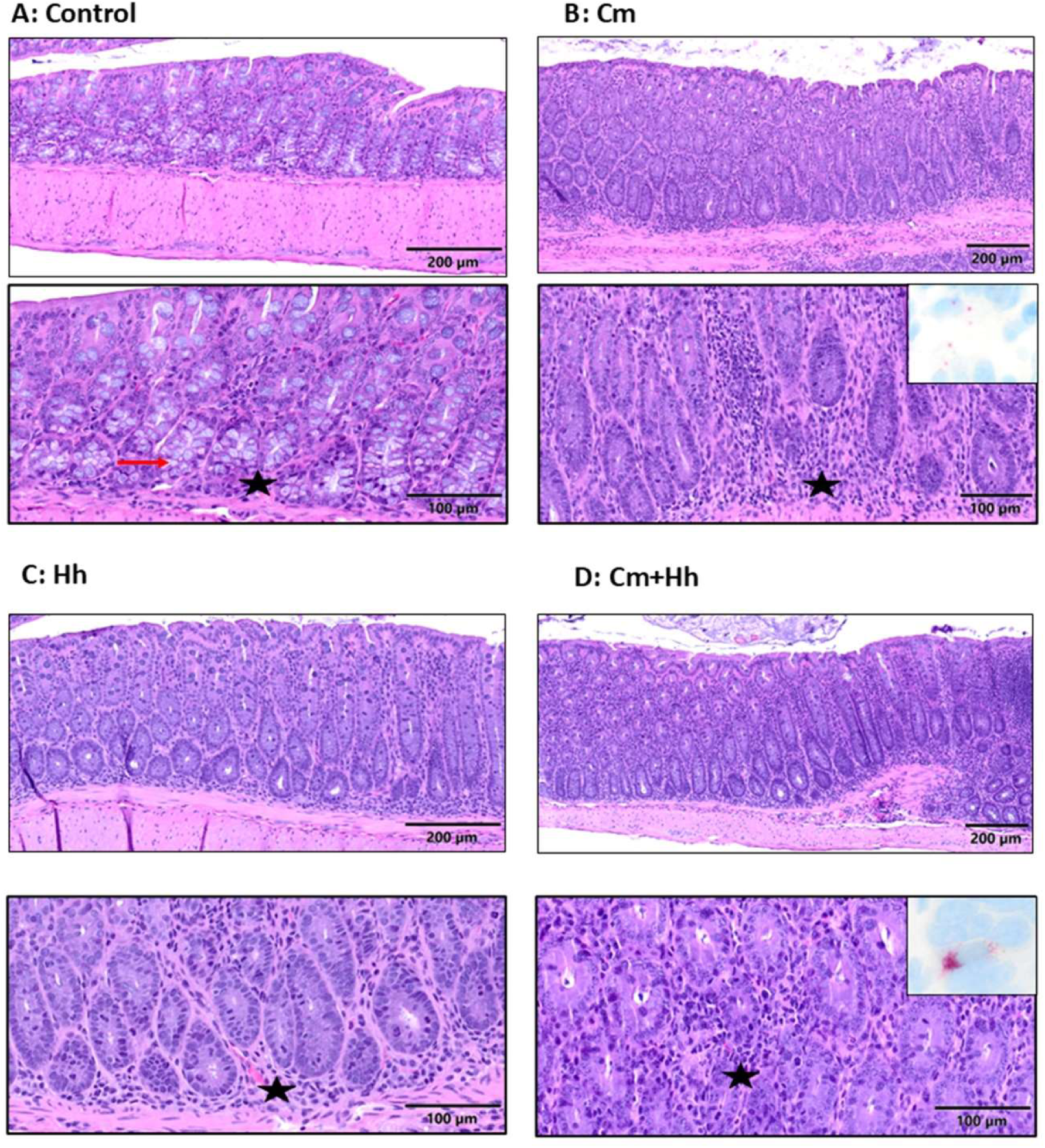
Colonic lesions in *Helicobacter hepaticus* (Hh) infected and free *Il*10^-/-^ mice coinfected with *Chlamydia muridarum* (Cm). Representative images from: (A) Colon of a Cm- and Hh-free *Il*10^⁻/⁻^ mouse showing occasional lymphocytes in the lamina propria (black star) and abundant goblet cells (red arrow; composite score = 2). (B) Colon from a Cm-infected *Il*10^⁻/⁻^ mouse with moderate colitis (black star), mild mucosal hyperplasia and moderate goblet cell atrophy (composite score = 11). Inset reveals Cm ISH signal (red) in mucosal epithelium. (C) Colon from an Hh-infected, Cm-free *Il*10^⁻/⁻^ mouse showing moderate to marked colitis (black star) with mild to moderate mucosal hyperplasia and moderate goblet cell atrophy (composite score = 11.5). (D) Colon from a Cm and Hh co-infected *Il*10^⁻/⁻^ mouse with epithelial hyperplasia and colitis (black star) and moderate goblet cell atrophy (composite score = 10). Inset reveals Cm ISH signal (red) in mucosal epithelium. Hematoxylin and eosin stain (10x and 20x magnification; insets, 40x magnification). Cm = *Chlamydia muridarum*; Hh = *Helicobacter hepaticus*; Control = Cm and Hh free; Cm + Hh = Confected with both bacterial species.

The individual pathology component scores as well as the composite scores for both the cecum and colon are provided in Figure 5; representative images demonstrating select pathologic changes are provided in Figure 4. While infecting *Il*10*^-/-^* mice with Cm did not impact the composite pathology score in the cecum, it did result in significant colonic pathology equivalent in magnitude to that resulting from Hh infection in this model. Hh caused significant pathology in both the cecum and colon. Infection of *Il*10*^-/-^* mice with both bacterial species did not increase the composite colonic pathology score more than the changes resulting from infection with either species alone.

**Figure 5.**
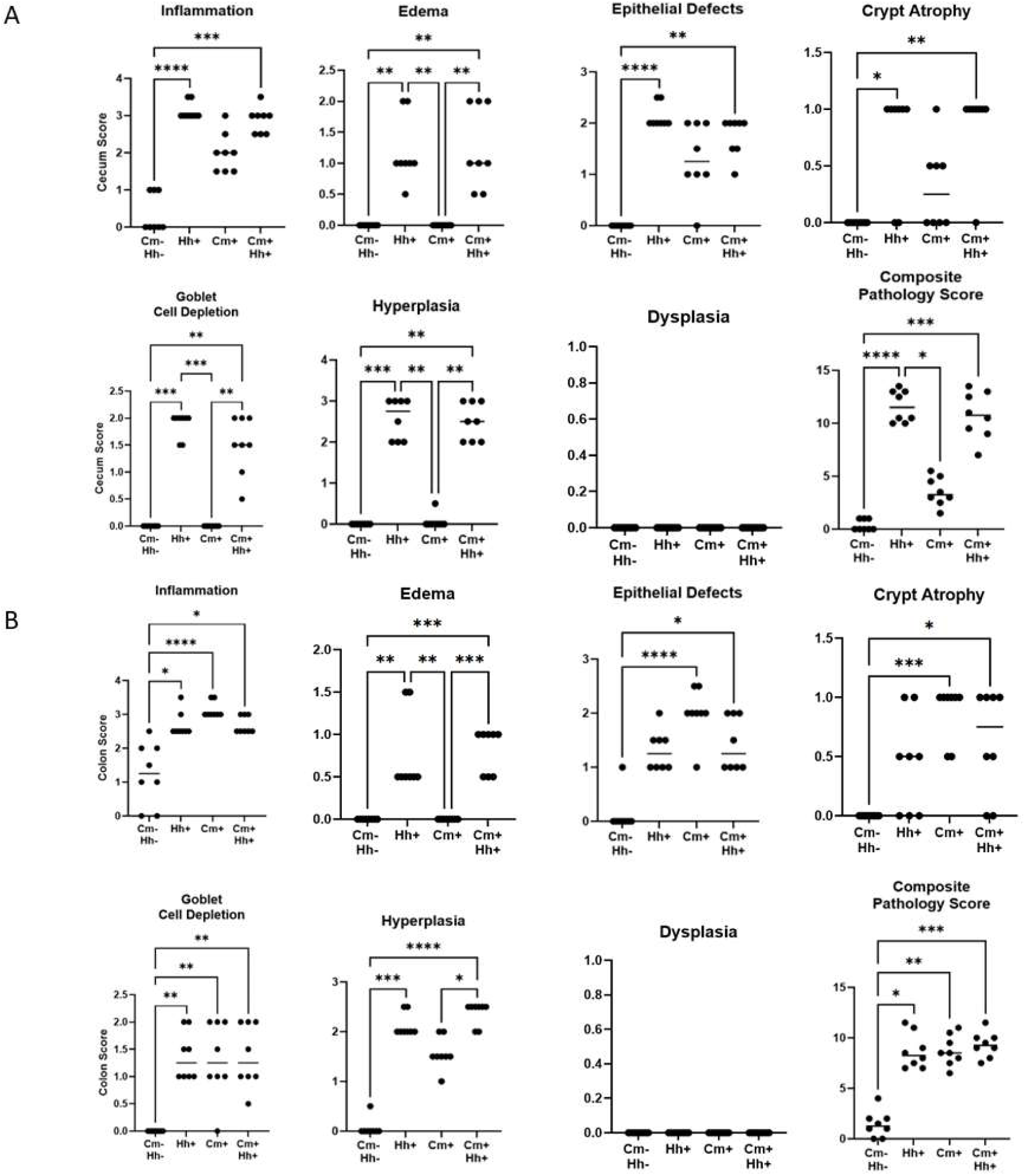
The effects of *Chlamydia muridarum* (Cm) on colitis in *Helicobacter hepaticus* (Hh)-infected and -free *Il*10^-/-^ mice. Pathology scores of the (A) cecum and (B) colon of female *Il*10^⁻/⁻^ mice (control), infected with Hh, infected with Cm, or co-infected with Cm and Hh (n=8 mice/group). Scores for inflammation, edema, epithelial defects, crypt atrophy, goblet cell depletion, hyperplasia, dysplasia and composite scores of all score parameters. The crossbars on the graphs indicate the median lesion scores for each parameter. *, P≤0.05; **, P≤0.01; ***; P≤0.001; ****, P≤0.0001

The composite score difference between Cm-free and -infected mice in the colon was a result of a significant increase in inflammation, epithelial defects, crypt atrophy, and goblet cell depletion following Cm infection. While epithelial hyperplasia was observed in the *Il*10*^-/-^* mice following Cm infection, the change was not significant. Crypt atrophy was also a principal component of the pathologic changes resulting from Cm infection, regardless as to whether the *Il*10*^-/-^* were infected with Hh. Crypt atrophy was inconsistently observed in *Il*10*^-/-^* mice infected with only Hh. While Hh infection of *Il*10*^-/-^*mice resulted in edema, edema was not a component of the pathology associated with Cm infection in this model.

While Cm infection of the *Il*10*^-/-^* mouse led to a minimal increase in the cecal composite pathology score, these increases were principally a result of inflammation and epithelial defects; however, none of these differences were statistically significant. Cecal edema, epithelial hyperplasia and goblet cell hyperplasia were the major pathologic features that were more severe in the Hh-infected as compared to the Cm-infected *Il*10*^-/-^* mice.

### Apc^Min/+^ DSS model of intestinal neoplasia

Mice were confirmed Cm-free or Cm-infected at euthanasia by qPCR as expected. Cm MOMP immunolabeling was also detected in tumors and normal intestinal mucosa (data not shown). None of the mice developed signs of illness for the duration of the experiment. As expected, DSS exposure resulted in increased tumorigenesis in the colon. Tumors were identified grossly as raised mucosal nodules (Figure 6B). Cm-infected Apc^Min/+^ mice had statistically fewer identifiable colonic tumors as compared to Cm-free Apc^Min/+^ mice (Figure 6A). The number of small intestinal and cecal tumors were not significantly different between Cm-free and Cm-infected Apc^Min/+^ mice. Overall tumor burden per mouse was reduced in the presence of Cm. All Cm-free Apc^Min/+^ mice exposed to 2% DSS developed gross intestinal tumors, however 1 of the Cm-infected mice failed to develop a colonic tumor. Two and 4 Cm-free mice failed to develop small intestinal and cecal tumors, respectively, as compared to the Cm-infected mice where 1 mouse failed to develop any small intestinal tumors and none of the 8 mice developed cecal neoplasia. All tumors evaluated histologically were classified as adenomas (Figure 6C).

**Figure 6:**
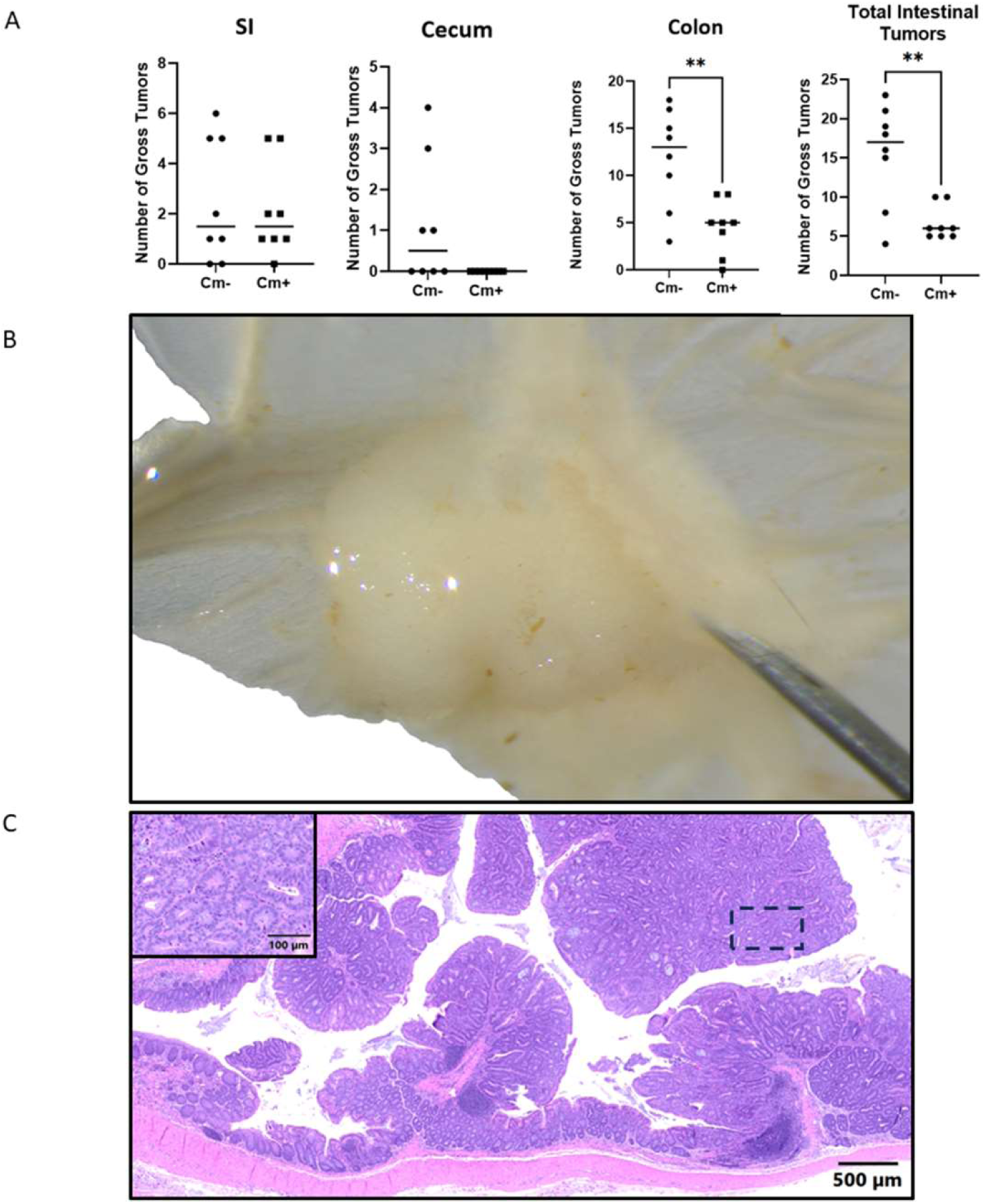
Colonic tumors in the small intestine, cecum and colon of Cm-infected and -free DSS treated, Apc^Min/+^ mice. (A) Gross tumor count in the small intestine, colon, cecum and total tumor count per mouse at necropsy of female control Cm-free Apc^Min/+^ mice or Cm-infected Apc^Min/+^ mice at 4-weeks post 2% DSS-treatment (n=8/group). The crossbar represents the median lesion score for each parameter. **p<0.005. (B) Representative photo of a gross tumor in the colon of Apc^Min/+^ mice 4-weeks post 2% DSS treatment at necropsy. (C) Representative H&E-stained images of a colonic tumor from Apc^Min/+^ mice 4-weeks post 2% DSS treatment. Tumors were characterized as adenomas (2x magnification, inset at 20x magnification).

## Discussion

Biosecurity is an integral and essential component of contemporary research mouse colony management as it is well recognized that there is a plethora of infectious agents that, while they may not cause clinical disease and/or pathology, can nevertheless confound research in which infected mice are utilized. It is for these reasons that most institutions maintain their rodent colonies, especially mice, in barrier facilities implementing specific husbandry and operational processes focused on keeping out a host of excluded infectious agents which may adversely impact the research for which the animals are used. Ascertaining which agents should be excluded is becoming ever more challenging as novel agents are identified, and the impact of the microbiota and agents which had historically been considered commensals is continuing to evolve.

Our group recently discovered and reported on the reemergence and surprisingly high prevalence of the bacterium Cm in research mouse colonies, and showed that Cm causes silent, but persistent infection of the large intestine of various commonly used murine strains and stocks.^1,6^ We further confirmed alterations of innate and adaptive immune responses following exposure by its natural route of infection.^6^ These findings lead to the question as to whether Cm can have a significant impact on the research in which infected mice are utilized, particularly those analyzing the intestine. A recent study revealed that Cm altered responses in the commonly utilized murine DSS-induced colitis model of inflammatory bowel disease. Cm infected DSS-treated mice had less pathology through an IL-22 dependent pathway.^7^ Therefore, we speculated that Cm would have a significant impact on other mouse models used to study gastrointestinal disease. We elected to assess Cm’s impact on 4 commonly used murine models, 3 of which are used to interrogate intestinal inflammation; infection with the bacterium Cr and the parasite Tm as well as the genetically engineered IL-10 knockout mouse model in which gut inflammation develops spontaneously in association with specific intestinal consortia. The DSS-treated Apc^Min/+^ mouse model of intestinal neoplasia was also assessed. While the study’s aim was not to determine the mechanism(s) by which Cm, if shown to influence any or all of these models, altered the models, but rather to aid us and others in determining the significance of its presence and assess whether the impact is sufficient to consider exclusion from some or all murine mouse colonies. These models were selected on their frequency of use as well as the likelihood, based on Cm’s pathophysiology, to alter the model phenotype.

The Cr mouse model of bacterial infection is commonly used to enhance the understanding of bacterial pathogenesis and mucosal immunity especially as it related to enterohemorrhagic and enteropathogenic *Escherichia coli* infection.^8^ Infection of B6 mice with 10^8^ - 10^9^ CFU of Cr results in transient colonic crypt hyperplasia typically lasting 2-3 weeks.^8^ Initially, Cr colonizes the colon and rapidly expands after infection. By day 7 post-infection, the bacterial load plateaus and by day 10, the infection starts to clear with complete clearance observed by 2-3 weeks.^8^ Hallmark histological changes associated with the developing colitis is the colonic crypt hyperplasia.^8^ We demonstrated that coinfecting Cm-infected mice with Cr significantly decreased the severity of colitis when compared to Cm-free mice inoculated with the same Cr dose. Cm and Cr coinfected mice had significantly less inflammation and goblet cell depletion, and while the magnitude of the epithelial hyperplasia and cell defects was not statistically different, both these parameters were also reduced in coinfected mice. We speculate that local and systemic upregulation of IFNγ, IL-17, and IL-22 following Cm infection resulted in the reduction of Cr-induced colitis.^6^ Th1, Th17 and Th22 and their respective cytokines, IFNγ, IL-17A, and IL-22 respectively, mediate host response to Cr infection.^15^ IFNγ deficient mice fail to clear Cr as a consequence of reduced activation of Cr-antigen specific T cells and the subsequent reduction in antibacterial IgG production, and macrophage phagocytosis.^16^ IL-17 and IL-22, in particular, are essential for protection against Cr.^8^ The latter is protective by upregulating epithelial repair and production of antimicrobial peptides.^15^ We excluded a single Cm and Cr coinfected mouse from the study, which was unexpectantly found dead 10 days after Cr inoculation with unusual and unexpected lesions.

We also examined the impact of Cm on the Tm induced colitis model in B6 mice. This model of human whipworm infection is useful for dissecting host-nematode interactions.^17^ A low dose of 50 ova elicits a Th1 response in which B6 mice become persistently infected and develop a chronic colitis, while mice gavaged with a high dose of 200 ova develop a Th2 response resulting in worm expulsion within 21 days.^17,18^ We elected to evaluate Cm’s impact on the latter model as we hypothesized the Th1 response and increase in IFNγ elicited by a preexisting Cm infection would increase susceptibility to Tm.^6^ However, there were no differences detected. All pathology criteria scored as well as the composite score were similar in Cm-free as compared to Cm-infected mice. The Th1 cytokines IL-12, IL-18 and IFNγ induce susceptibility to Tm infection.^17^ IFNγ is thought to be the principal mediator as depletion results in expulsion of Tm in susceptible strains.^19^ IL-18 and IL-12 induce IFNγ production.^20^ *Il*12b^-/-^ and *Il*18^-/-^ animals are resistant to infection.^21^ Resistance in *Il*12^-/-^ mice is primarily mediated by reducing IFNγ production; however, resistance in *Il*18^-/-^ is a result of IL-13 suppression inhibiting the Th2 response.^21^ Administration of recombinant IL-12 or IL-18 to a resistant strain results in susceptibility to infection, further implicating the Th1 cell response is key to susceptibility to chronic Tm infection.^21^ The observed lack of differences in our study may reflect Cm’s other effects. Although Cm infection elicits an increase in IFNγ, it also increases IL-22 and IL-17.^6^ IL-22 deficiency leads to susceptibility to Tm infection in normally resistant mice.^22^ Loss of IL-22 decreases goblet cell proliferation and mucous production impairing worm expulsion.^22^ Therefore, even though Cm infection leads to an increase in production of IFNγ, the increase in IL-22 may have counteracted its impact. Given these results, it would be interesting to evaluate Cm’s impact on the low dose B6 model as the heightened Th1 cell response could increase pathology and worm burden.

Cm’s impact on the *Il*10^-/-^ model of gut inflammation was also examined as it has a clinicopathologic phenotype influenced by the microbiota.^23–25^ The loss of IL-10 leads to colitis as a result of an aberrant immune response in the presence of specific agents such as Hh.^23^ We demonstrated that Cm induced colitis of similar magnitude to that caused by Hh. Therefore, Cm can be added to the list of agents capable of inducing colitis in the model. Interestingly, coinfection with both agents did not lead to colitis of greater severity than either bacterium caused alone. This finding may reflect that both bacteria induce colitis by the same mechanism.

IFNγ, IL-17 and IL-22 are upregulated in the absence of IL-10.^23,26^ IFNγ has historically been linked to the development of colitis in Hh-infected *Il*10^-/-^ mice as intestinal inflammation is reduced following administration of neutralizing monoclonal antibodies.^27^ Recent studies have shown that mice deficient in both IFNγ and IL10^-^still developed colitis following Hh infection suggesting that colitis is likely induced by other pathways.^28^ Although IL-17 was linked to colitis in this model, there are more recent reports of mice deficient in both IL-17 and IL-10 developing more severe colitis than mice deficient only in IL-10.^27^ IL-22 is also overexpressed in *Il*10^-/-^ mice and is associated with the colitis phenotype.^26^ *Helicobacter* spp. infected mice deficient in both IL-22 and IL-10 remained colitis free suggesting that IL-22 is a major driver of colitis in this model.^26^ Given that IFNγ, IL-17and IL-22 are upregulated following Cm infection, it is not surprising that Cm is sufficient to induce colitis in *Il*10^-/-^ mice in the absence of Hh.^6^

Interestingly, 2 of 16 *Il*10^-/-^ mice were negative for Cm by qPCR at euthanasia despite being qPCR positive following cohousing 70 days prior. One of these 2 mice was coinfected with Hh while the other was Hh-free. Both were singly housed after cohousing which could have contributed to a reduction in Cm burden to undetectable levels by qPCR as they would not have been continuously exposed to Cm by a Cm-shedding cage mate. Despite negative qPCR results, intestinal pathology was similar to other Cm-positive animals. Interestingly, all Cm-infected *Il*10^-/-^ were also MOMP IHC negative, which was not observed in any of the other models we evaluated, although Cm was detectable in most of the *Il*10^-/-^ mice evaluated via ISH. Of the 2 mice negative for Cm by qPCR, 1 was also negative by ISH. Collectively, these findings indicate that the Cm burden in the large intestine decreases temporally in this model with only trace amounts of Cm nucleic acid detectable in the colonic mucosa by the end of the experimental period. Given that IL-10 is a potent anti-inflammatory cytokine that modulates immune responses, it is plausible that the exaggerated inflammatory response led to the clearance of most chlamydial inclusions from the intestines. This notion is supported by prior studies utilizing *Il*10^⁻/⁻^ mouse models, which demonstrated a progressive reduction in *Chlamydia trachomatis* burden in the lungs beginning 7 days post-infection.^29^ Although proinflammatory cytokine levels were not assessed in the colonic mucosa of Cm-infected *Il*10^⁻/⁻^ mice, it is well established that upregulation of IFNγ production by epithelial cells plays a critical role in host defense by depleting intracellular tryptophan, an essential nutrient for *Chlamydia* replication within host tissues.^30^ Future studies should elucidate the immunoregulatory pathways influenced by Cm in the colon of *Il*10^⁻/⁻^ mice.

Finally, we assessed the impact of Cm infection on the DSS-treated Apc^Min/+^ mouse of colonic neoplasia. Importantly, Cm infection significantly decreased tumor burden in the model. We elected to use the DSS-treated model as DSS exposure stimulates colonic neoplasia rather than the small intestinal neoplasia that predominates in untreated Apc^Min/+^ mice.^31,32^ As Cm infection preferentially colonizes the large intestine, we speculated that it would be more likely to influence the DSS-treated model.^6^ Although the specific mechanism by which DSS impacts colonic tumorigenesis is unknown, the induction of colonic neoplasia is likely influenced by the resulting colitis.^31^ Our findings are not surprising considering Cm alleviated DSS-induced colitis by stimulating IL-22 production and the cytokine has been shown to be protective in chemically-induced models of tumorigenesis by decreasing inflammation.^33,21,7^ Interestingly, IL-22 can also be tumorigenic as *Il*22^-/-^Apc^Min/+^ mice develop fewer tumors as compared to *Il*22^+/+^Apc^Min/+^ mice.^33,34^ The mechanism is believed to be a result of the epithelial turnover induced by IL-22 suggesting that multiple pathways could impact tumorigenesis in Cm-infected, DSS-treated Apc^Min/+^ mice.^34^ It remains unknown whether Cm infection could influence neoplasia in untreated Apc^Min/+^ mice. Cr infection of Apc^Min/+^ mice resulted in increased colonic neoplasia; the mechanism was thought to result from increased epithelial turnover.^35^ Although Cm infection has not been reported to induce epithelial hyperplasia, we observed epithelial hypertrophy in Cm-infected *Il*10^-/-^ mice.^4^ Future research could examine the impact of Cm in the untreated Apc^Min/+^ mouse model.

Cm infection significantly impacted 3 of the 4 GI models evaluated in this study providing further and compelling evidence that Cm status should be considered when utilizing these models. One reason to consider excluding Cm is the sensitivity of available diagnostics for its’ detection and, more importantly, the relative ease by which Cm can be successfully eradicated from most mouse strains using antibiotic therapy.^36^ This contrasts with other prevalent murine infectious agents, such as MNV, that while ideally should be excluded, the process of establishing MNV-free colonies would be extremely challenging in that numerous unique strains of mice present in many colonies would need to be rederived. Careful consideration should also be given to using previously Cm-infected, antibiotic-treated mice as research subjects as the impact of prior Cm infection is unknown. Ideally offspring of treated mice should be utilized, although epigenetic multigenerational effects remain possible.

## Acknowledgements

We thank Jacqueline “Jackie” Candelier, Sockie Jiao and the staff of the Laboratory of Comparative Pathology for their assistance with necropsies, histology, immunohistochemistry and in-situ hybridization assays, Mohd Anees Ahmed for his assistance with *Citrobacter rodentium* culture and infection, and Mengze Lyu for his assistance with *Helicobacter hepaticus* culture.

## Abbreviations

B6: C57BL/6J
BCS: body condition score
C: BALB/cJ
Cm: *Chlamydia muridarum*
Cr: *Citrobacter rodentium*
Tm: *Trichuris muris*
Hh: *Helicobacter hepaticus*
DSS: dextran sodium sulfate
Apc: adenomatous polyposis coli
H&E: hematoxylin and eosin
IHC: immunohistochemistry
ISH: in situ hybridization
GI: gastrointestinal
GIT: gastrointestinal tract

